# Effects of priority on strain-level composition of the honey bee gut community

**DOI:** 10.1101/2025.04.21.649814

**Authors:** Korin Rex Jones, Yulin Song, Sabrina Sakurako Rinaldi, Nancy A Moran

**Author notes:** Address correspondence to Korin Rex Jones.

## Abstract

Host-associated microbiomes are complex communities shaped by interactions between members. The Type VI Secretion System (T6SS), among other bacterial weapons, allows gram-negative bacteria to deliver toxic effectors into competitors. In this study, we investigated the impact of differential colonization timing on the competitive advantage associated with T6SS possession using *Snodgrassella alvi*, a core symbiont of the honey bee gut microbiome. Following a timeline based on the natural establishment window of the gut microbiome, we sequentially inoculated newly emerged bees with fluorescently labeled strains that differed in presence of the T6SS-1. When inoculated simultaneously, the T6SS-1-possessing strain (wkB2) consistently excluded the T6SS-1-lacking strain (wkB332); however, when given a five-day advantage, the second strain was consistently excluded regardless of strain identities. With a one-day advantage, the effect of priority was weakened, but wkB332 was able to persist following introduction of wkB2. Utilizing a wkB2 T6SS-1 knockout strain, we repeated our 24-hour priority experiments and found that the T6SS-1 contributes to invasion outcomes along with other mechanisms of competition. Through fluorescent microscopy, we explored how coexisting strains in these experimental scenarios organize spatially within the bee ileum. Our results demonstrate that colonization timing can have lasting consequences for strain composition of the established microbiome. These findings illustrate the influence of stochastic processes in microbial community assembly and emphasize that differences in colonization timing may alter competitive outcomes between taxa, impacting taxon coexistence.

**Importance:** The bacterial gut communities of honey bees possess considerable strain-level diversity between hives, between individual bees and within individual bees. However, the factors underlying strain coexistence are unclear. Here, we provide support for timing of colonization, or priority effects, as one factor driving this strain level diversity. Our results show that priority inoculation can prevent colonization by subsequent competing bacterial strains and mitigate advantages conferred through bacterial weaponry. Further, a brief window of priority can facilitate the coexistence of strongly and weakly competitive strains within single bees. These results add to our understanding of the impacts of priority effects in host-associated microbial communities and provide support for the development of future probiotic strategies aimed directing microbiome structure to benefit honey bee health.

## Introduction

Host-associated microbiomes are complex communities that consist of interacting species of microorganisms. The ultimate structure of these communities, specifically which microbes are present and in what proportions, can have consequences for host health and development [1–4]. Microbiome composition has been shown to impact host obesity[5], disease susceptibility [6, 7], vulnerability to toxins [4, 8], and behavior [9, 10]. Because of this relationship between structure and function, understanding the processes governing how communities assemble continues to be an important goal in microbial ecology [11, 12].

Broadly, the assembly of ecological communities is understood to be governed by four ecological processes: selection, dispersal, speciation, and drift [13]. Among different systems and conditions, the relative influence of each of these factors will fluctuate [14]. Even among equivalent habitats, stochasticity may result in differences in microbiome community composition [15, 16]. For example, differences in the arrival order of bacterial colonists can sometimes impact microbiome composition, commonly described as priority effects [17–19]. Early arriving species may dominate resources (niche preemption) or modify the environment (niche modification) in a way that hinders the colonization ability of subsequent species [12, 20, 21]. In flower nectar, for example, early arriving *Acinetobacter* can lower pH, preventing future colonization by yeasts [21]. Because priority effects are expected to occur in situations where fitness differences between competing taxa are low and stabilizing forces are absent [22], they may be especially prominent in strain-strain competition during colonization.

In addition to stochastic processes, deterministic processes also influence microbiome structure [23, 24]. Bacteria can possess a diverse array of weaponry, providing advantages for certain strains over others [25, 26]. The Type VI Secretion System (T6SS) is one such mechanism, capable of delivering effector proteins into neighboring cells in a contact-dependent manner, with individual strains possessing immunity mechanisms against their own effectors [27]. Initially implicated in pathogenesis, the T6SS has since been found to determine competitive outcomes within and among bacterial species [28, 29]. For example, an in planta coinfection assay found that the soil bacterium *Agrobacterium tumefaciens* utilized the competitive advantage provided by its T6SS to attack and kill *Pseudomonas aeruginosa* cells[30]. Similarly, experimental inoculations in the *Vibrio*-squid system demonstrate that strains of *Vibrio fischeri* utilize the T6SS during strain-strain competition *in vivo* [31]. In T6SS-mediated antagonism, the more abundant strain enjoys an advantage, as it possesses immunity mechanisms against its own weaponry, resulting in positive feedback.

The gut microbiome of honey bees (*Apis mellifera*) is a simple, conserved community of bacteria largely dominated by a few key members (*Bartonella, Bifidobacterium, Bombilactobacillus* (formerly “Firm-4”), *Commensalibacter, Frischella, Gilliamella, Lactobacillus* nr. *melliventris* (formerly “Firm-5”), and *Snodgrassella*) [32, 33]. Prior research on this community suggests that both stochasticity in dispersal and deterministic competitive processes likely influence community composition [34–36]. Within the genera *Snodgrassella* and *Gilliamella*, which together form a densely packed biofilm in the ileum region [37], different approaches have shown strain-level variation among hive-mates that may be driven by stochastic processes such as priority effects [34, 38]. Both *Snodgrassella* and *Gilliamella* include strains that encode T6SSs and a diverse array of associated effectors [39, 40].

*S. alvi* is a primary colonist that characteristically forms a biofilm on the ileum wall, where it may interact with the host immune system and influence the colonization of other taxa [33, 41]. The *S. alvi* type strain wkB2 possesses two T6SSs. T6SS-1 facilitates intraspecific competition, and T6SS-2 promotes colonization of the host [35]. Other strains of this species can possess one, both, or neither of these T6SSs [39, 40]. It is unclear how *S. alvi* strains that lack immunity to T6SS-1 effectors might persist in populations of host bees. Stochasticity in colonization timing within individual bees, which facilitate priority effects, may be one explanation [34].

In the current study, we inoculated newly-emerged, microbiota-deprived bees with one of three fluorescently tagged [42] strains of *S. alvi* in a pair-wise sequence and included delays between the introduction of the first and second strain to better understand colonization dynamics in the honey bee gut. We chose two naturally occurring strains that differ in their encoded T6SS genes; wkB2 (T6SS-1 and T6SS-2) and wkB332 (T6SS-2 only). We also included a T6SS-1 knockout of wkB2 to disentangle the impact of T6SS antagonism from other contrasting strain characteristics. We hypothesized that if a strain vulnerable to T6SS attack were to be established prior to the introduction of a T6SS-possessing strain, the temporal advantage would outweigh its vulnerability and facilitate its persistence.

## Materials and Methods

### Bacterial Cultures

*S. alvi* strains wkB2 and wkB332 were previously isolated from guts of *A. mellifera* and were used in a previous study on strain interactions ([35]). For the wkB2 T6SS-1 knockout strain, we used wkB2 ΔtssE-1, which was constructed in a previous study ([35]).

Strains were cultured on Columbia agar supplemented with 5% defibrinated sheep blood (hereafter CBA). *Escherichia coli* cultures were grown on LB agar, LB broth, or CBA. Culture media was supplemented as needed with 0.3 mM diaminopimelic acid (DAP), 12.5 μg/mL tetracycline (tet), 50 μg/mL kanamycin (kan), and/or 60 μg/mL spectinomycin (spec). Plated cultures were grown in a 5% CO^2^ incubator at 35°C.

### Strain Transformation

All strains used in this study were transformed using the Pathfinder plasmid system described in Elston et al. [42]. Briefly, cultures of E. coli MFDpir containing GFP (pSL1-GFP) or RCP (red chromoprotein; pSL1) plasmids were started on LB agar supplemented with kan and DAP. Cultures of *S. alvi* were simultaneously started on CBA. The next day, single colonies from the MFDpir strains were placed in LB broth supplemented with kan and DAP overnight. The following day, *S. alvi* strains were scraped and suspended in 500 μL PBS. *S. alvi* suspensions and 500 μL of MFDpir strains were spun down at 14000 rpm for three minutes. The liquid was replaced with a fresh 500 μL aliquot of PBS and vortexed to resuspend the pellet. After repeating this a second time, the OD of each suspension was measured, and strains were mixed at a ratio of 1:10 donor to recipient. 100 μL of each mixture was spot plated on CBA plates supplemented with DAP and placed in an incubator overnight. The following day, cultures were scraped and resuspended in 500 μL PBS. Full and 1:10 dilutions of these resuspended cultures were then plated onto kan/tet or kan/tet/spec supplemented CBA plates. Single fluorescent colonies were then picked and plated on kan/tet CBA plates. GFP and RCP variants for each strain were created and stored in 25% glycerol at -80^o^C.

### Inoculum Preparation

Stock cultures of *S. alvi* strains were plated on kan CBA plates and incubated for 2 days. Single colonies were then picked from these plates, struck onto new kan CBA plates and incubated for two days. Plates were then scraped and resuspended in a 1.5 mL microcentrifuge tube containing 500 μL PBS. Opitcal densities of these suspensions were measured in a BioSpectrometer (Eppendorf, CT), and the volume of culture needed to form 1 OD of solution was placed in a new microcentrifuge tube and spun down at 14,000 rpm to separate the bacterial pellet from the supernatant. After centrifugation, the supernatant was discarded. Bacteria were resuspended in 500 μL PBS and 500 μL 1:1 sterile sucrose:water solution. The sugar/water solution was added at the time of hand feeding to minimize potential cell death caused by the solution.

### Priority Inoculation Experiments

To obtain microbiota-deprived newly emerged workers of *A. mellifera*, we followed an established protocol (see Motta et al. 2024 [35]) with minor modifications. Briefly, brood frames were collected from two hives located on the University of Texas at Austin campus. Pupae in late development, characterized based on body and eye pigmentation, were extracted from frames using sterile forceps. Collected pupae were then placed into sterilized plastic containers lined with Kimwipes and supplied with sterile 1:1 sucrose:water solution. Pupae were kept in a 35°C incubator with ∼60% relative humidity to mimic hive conditions. After a few days, we transferred newly emerged, microbiota-deprived workers to sterile feeding tubes (0.5 mL microcentrifuge tubes with the ends cut off) (Figure 1A). Bees were starved for ∼2 hours and then hand-fed a 5 μL treatment inoculum. This inoculum contained a sucrose solution composed of either a single strain, or an evenly proportioned mixture of two strains. After feeding, bees were placed into individual polypropylene petri dishes (Eisco, NY) containing a 0.5 mL microcentrifuge tube filled with sterile 1:1 sucrose:water solution. After the priority period had elapsed for the initial strain, bees were briefly chilled to restrict movement and then placed in new sterile feeding tubes and starved for ∼2 hours, before being fed the second inoculum and returned to their individual petri dishes. Bees were kept in the incubator at 35°C until the end of the experiment.

**Figure 1:**
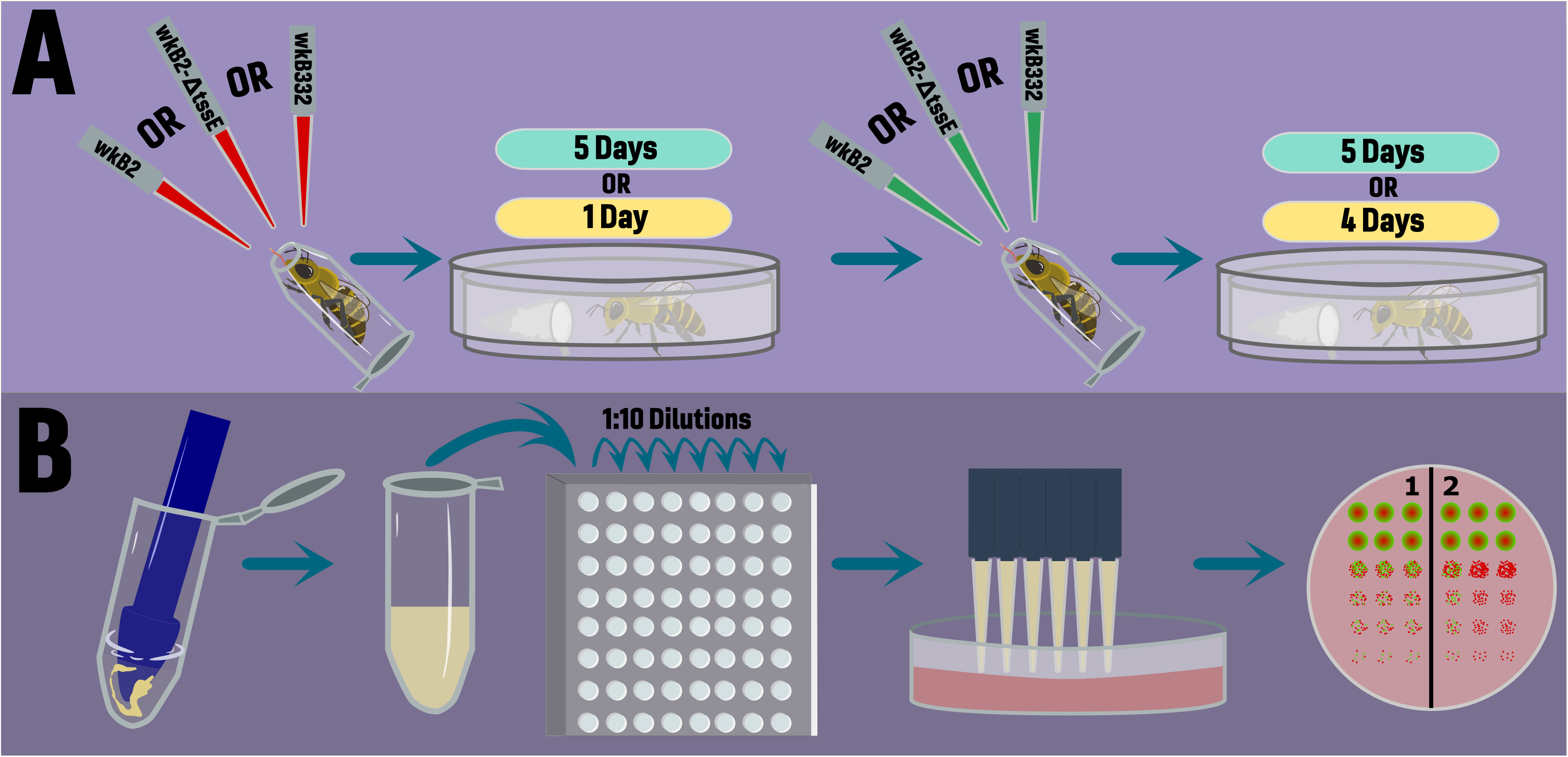
Concept diagram explaining experimental design. **A)** Newly emerged, microbiota-deprived honey bees were sequentially inoculated with fluorescently tagged strains of *S. alvi* (wild type wkB2, wkB2 ΔtssE-1knockout, or wild type wkB332) over two experimental timelines. Bees were either inoculated with a secondary strain after 5 days, followed by 5 days of incubation or after 1 day, followed by 4 days of inoculation. **B)** Once the experiment was completed, bees were dissected, their guts were homogenized, and dilution series were conducted in 96-well plates. These dilutions were then plated on Columbia blood agar plates and CFUs were identified and counted based on fluorescence.

To assess colonization outcomes at the end of the incubation period, bees were chilled and their guts dissected with flame-sterilized forceps. Extracted guts were homogenized in 100 μL PBS using a BioVortexer (BioSpec, OK) and sterilized polypropylene pestles for 30 s (Figure 1B). Pestles were then rinsed with 400 μL of PBS, bringing the total volume to 500 μL, and stored on ice. 20 μL of each sample was then used to prepare 10-fold serial dilutions. 10 μL of each dilution was plated in triplicate on kan-supplemented CBA plates and incubated for three days. Gut homogenates for microbiota-deprived bees were plated on both CBA and kan supplemented CBA. Colony-forming units (CFUs) were then counted for each sample from the dilution with the largest number of countable colonies (distinguishable individual colonies).

### Individual Experimental Design

Our experimental design consisted of two types of priority inoculation experiments (Figure 1A). First, we inoculated newly emerged, microbiota-deprived bees with a single strain of *S. alvi*, allowed five days for that strain to colonize, introduced the second strain and then waited an additional five days before concluding the experiment and performing dissections for CFU counts (5d/5d). The normal timeline for establishment of the honey bee microbiome is ∼4 days[43]; therefore, this experimental design gives insight into a strain attempting to invade a “fully colonized” ileum. Second, we employed a design in which we restricted the initial strain to only one day of priority before the second strain was introduced and then four additional days for the bacteria to colonize the host bee (1d/4d) (Figure 1A). We also included a mixed treatment within this design in which strains were introduced simultaneously on day one. These samples were dissected on the same day as the other bees in the 1d/4d experiment. With this strategy we sought to more closely mimic natural scenarios that may take place in the hive environment by keeping our timeline within the natural period of microbiome establishment [43]. We performed four experiments sequentially: 1) 5d/5d with both fluorescent markers for each wild type (WT) strain, 2) 1d/4d with both fluorescent markers for each WT strain, 3) 1d/4d with single fluorescent markers for our two WT and T6SS knockout strains, 4) 1d/4d with alternate fluorescent markers for our two WT strains and T6SS knockout strain. Experiments 1 and 2 used bees from a single hive and are analyzed individually. Experiments 3 and 4 consisted of bees from a second hive and are combined into a single dataset. All data used in this study are available in the supplementary material.

### Imaging honey bee ileum with fluorescent microscopy

Honey bees were raised and inoculated as above. Three bees from each treatment were selected and processed for imaging. Whole guts were dissected and mounted with PBS onto glass slides under a Leica MDG41 stereoscope and Malpighian tubules were separated from view of the ileum. The ileums of dissected guts were then imaged using Nikon NIS-Elements software (AR 5.30.05 Build 1559) and a Nikon Eclipse TE2000-U inverted fluorescent microscope in GFP and RCP channels to visualize bacterial colonization.

### Statistical analysis

Statistical analyses were performed in the R v4.3.1 environment [44] via R studio [45](build 524).

To understand if the identity of the initial strain impacted either the ability of the secondary strain to invade or the relative abundance of the secondary strain, we performed Kruskal-Wallis rank sum tests. To understand if strain inoculation order influenced invasion success of secondary strains or impacted strain relative abundance, we performed pairwise Wilcoxon rank sum tests. P values from our Wilcoxon rank sum tests were corrected for multiple comparisons using the Benjamini Hochberg method [46]. Plots were created using the ggplot (3.4.2)[47], ggpubr (v0.6.0)[48], scales (v1.2.1)[49], and phyloseq (1.44.0)[50] packages.

## Results

Experimentally inoculating microbiota-deprived, newly emerged honey bees with one strain of *S. alvi* and a second strain five days later led to strong priority effects (Figure 2). Samples were dominated by the initial strain, with no treatment having a mean initial strain percentage lower than 87%. The identity of the initial strain (wkB2 or wkB332) did not correspond with a difference in whether the second strain was able to invade (KW chi-squared = 1.8, df = 1, p-value = 0.19)(Table 1). Similarly, there was no observable difference in the percentage of the secondary strain based on initial strain identity (KW chi-squared = 1.5, df = 1, p-value = 0.22) (Figure 2A).

**Table 1.** Ability of strains to invade in experimental inoculations.

| 5 Day/5 Day Experiment |  |  |  |  |
| --- | --- | --- | --- | --- |
|  | Inoculation Order | Invaded* | Not Invaded | Total |
|  | wkB2>wkB332 | 1 | 17 | 18 |
|  | wkB332>wkB2 | 3 | 11 | 14 |
| 1 Day/4 Day Experiment |  |  |  |  |
|  | Inoculation Order | Invaded | Not Invaded | Total |
|  | wkB2>wkB332 | 5 | 22 | 27 |
|  | wkB332>wkB2 | 22 | 14 | 36 |

\*Invaded means that the second strain was present at any level at the time of sampling.

**Figure 2:** Relative abundance plots based on CFU counts of *S. alvi* indicating proportions of wild type wkB2 and wkB332 present during **A)** 5 day/5 day or **B)** 1 day/4 day experimental timelines based on inoculation order.

Restructuring our inoculation timeline to be within the normal window of microbiome establishment in honey bees, we examined priority effects when a strain is given only one day of priority. We first performed a 1d/4d experiment and found that invasion success of the second strain depended on initial strain identity, with wkB2 being invaded less often (KW chi-squared = 11.2, df = 1, p-value = 0.0008) (Table 1). Percentages of the secondary strain also were lower when wkB2 was the initial strain (KW chi-squared = 13.6, df = 1, p-value = 0.0002)(Figure 2B).

To understand the potential role of the Type VI Secretion System in these colonization outcomes, we performed another 1d/4d experiment in which we included a T6SS-1 knockout mutant (wkB2 ΔtssE-1). When given priority, wkB2 WT was far more successful at preventing invasion by wkB332 than was the T6SS-1 knockout strain (Pairwise-Wilcoxon Rank Test p=0.0008)(Table 2). Similarly, wkB332 was invaded by wkB2 WT more often than by the wkB2 ΔtssE-1knockout strain (Pairwise-Wilcoxon Rank Test p=0.032)(Table 2).

**Table 2.** Ability of strains to invade in experimental inoculations in 1d/4d experiment that includes wkB2 with inactivated T6SS-1.

| Inoculation Order | Invaded* | Not Invaded | Total |
| --- | --- | --- | --- |
| wkB2>wkB332 | 0 | 33 | 33 |
| ΔtssE>wkB332 | 9 | 19 | 28 |
| wkB332>wkB2 | 34 | 4 | 38 |
| wkB332>ΔtssE | 26 | 12 | 38 |

\*Invaded means that the second strain was present at any level at the time of sampling.

We next evaluated the overall impact of priority versus simultaneous inoculation on proportional abundances of wkB332 when paired with either wkB2 WT or wkB2-ΔtssE-1. For wild type pairings, wkB2 WT enjoyed a greater advantage when it had a 1d priority versus when introduced simultaneously with wkB332 (Pairwise-Wilcoxon Rank Test p=0.049). However, only 3 of the 23 simultaneously inoculated communities contained wkB332(Figure 3). For wkB332, we also saw an increase in relative abundance in priority treatment samples compared to simultaneous inoculations when the second strain was the wkB2 WT (Pairwise-Wilcoxon Rank Test p=7.8e-07) (Figure 3). In pairings with the T6SS knockout strain, introducing wkB2-ΔtssE-1 first also led to an increase in its relative abundance compared to samples from simultaneous treatments (Pairwise-Wilcoxon Rank Test p=1.2e-05) (Figure 3). For wkB332, however, we did not see an increase in relative abundance compared to simultaneous inoculations when the second strain was wkB2-ΔtssE-1 (Pairwise-Wilcoxon Rank Test p=0.45) (Figure 3).

**Figure 3:**
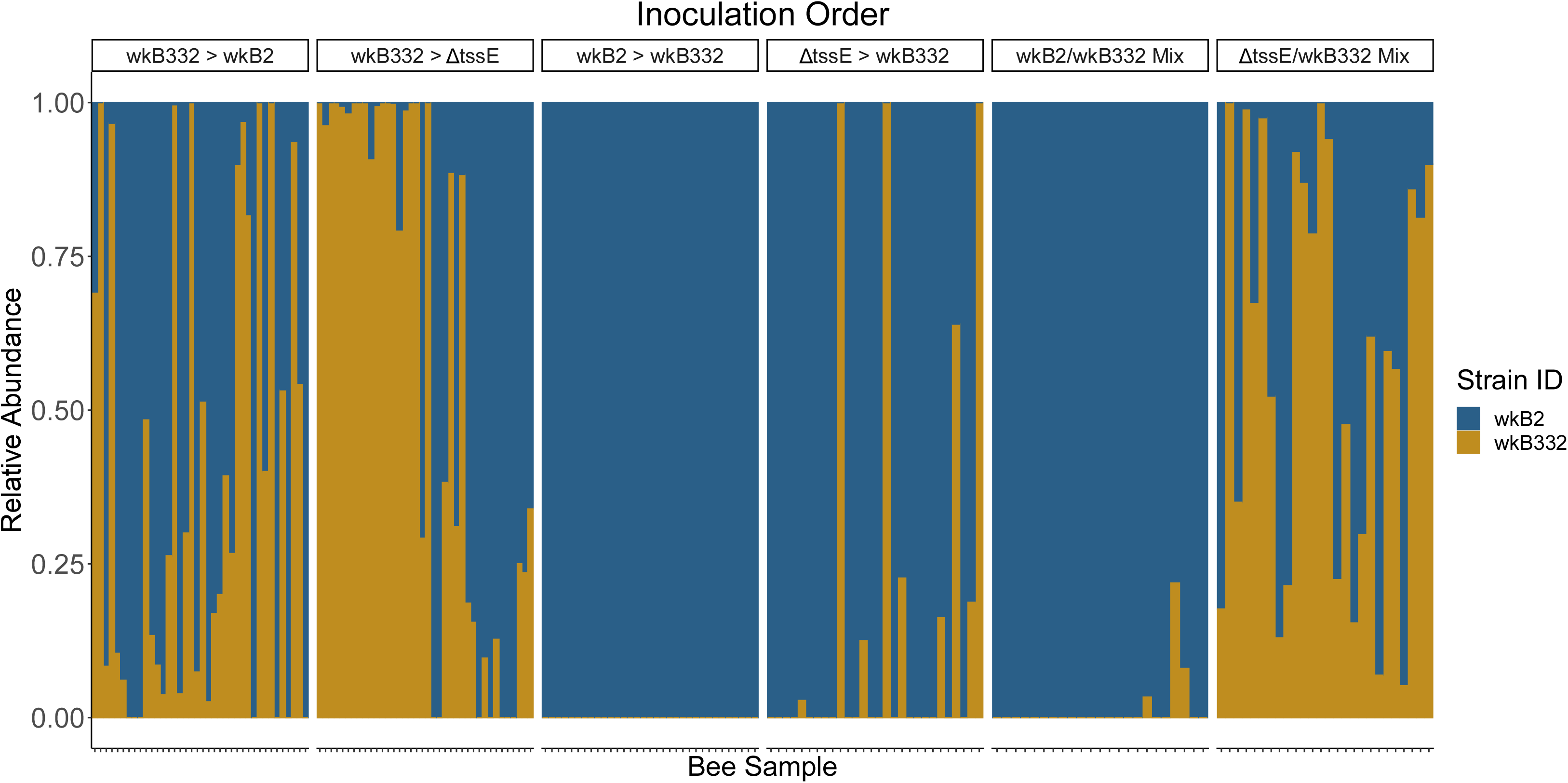
Relative abundance plot based on CFU counts of *S. alvi* indicating proportions of wild type wkB2, wkB2 ΔtssE-1, and wild type wkB332 based on experimental inoculation order.

To understand the potential influence of the T6SS on competitive outcomes between our isolates during our inoculation experiments, we analyzed differences in abundances of wkB332 between priority treatments. Compared to samples from treatments in which wkB2 WT was introduced first, wkB332 showed a higher relative abundance when wkB2-ΔtssE-1 was given priority (Pairwise-Wilcoxon Rank Test p=0.0009) (Figure 3). When wkB332 was first, wkB332 had a greater advantage over wkB2-ΔtssE-1 than over wkB2 WT(Pairwise-Wilcoxon Rank Test p=0.048) (Figure 3).

To better understand scenarios that implied coexistence between strains, we examined the spatial context of these outcomes by imaging intact ileums from a subset of bees from each treatment using fluorescence microscopy. We observed a general preference of all strains for the anterior end of the ileum, with the densest fluorescence observed near the pylorus (Figure 4). When wkB332 was the priority inoculum, we saw large, continuous patches of colonization adorned with considerably smaller patches of wkB2/ wkB2-ΔtssE-1 at both the anterior and posterior ends of the ileum (Figure 4A,B). When wkB2-ΔtssE-1 was introduced first, we saw an overlap in fluorescent signal between strains (Figure 4C). We also observed overlap between wkB332 and wkB2-ΔtssE-1 when inoculated simultaneously (Figure 4D).

**Figure 4:**
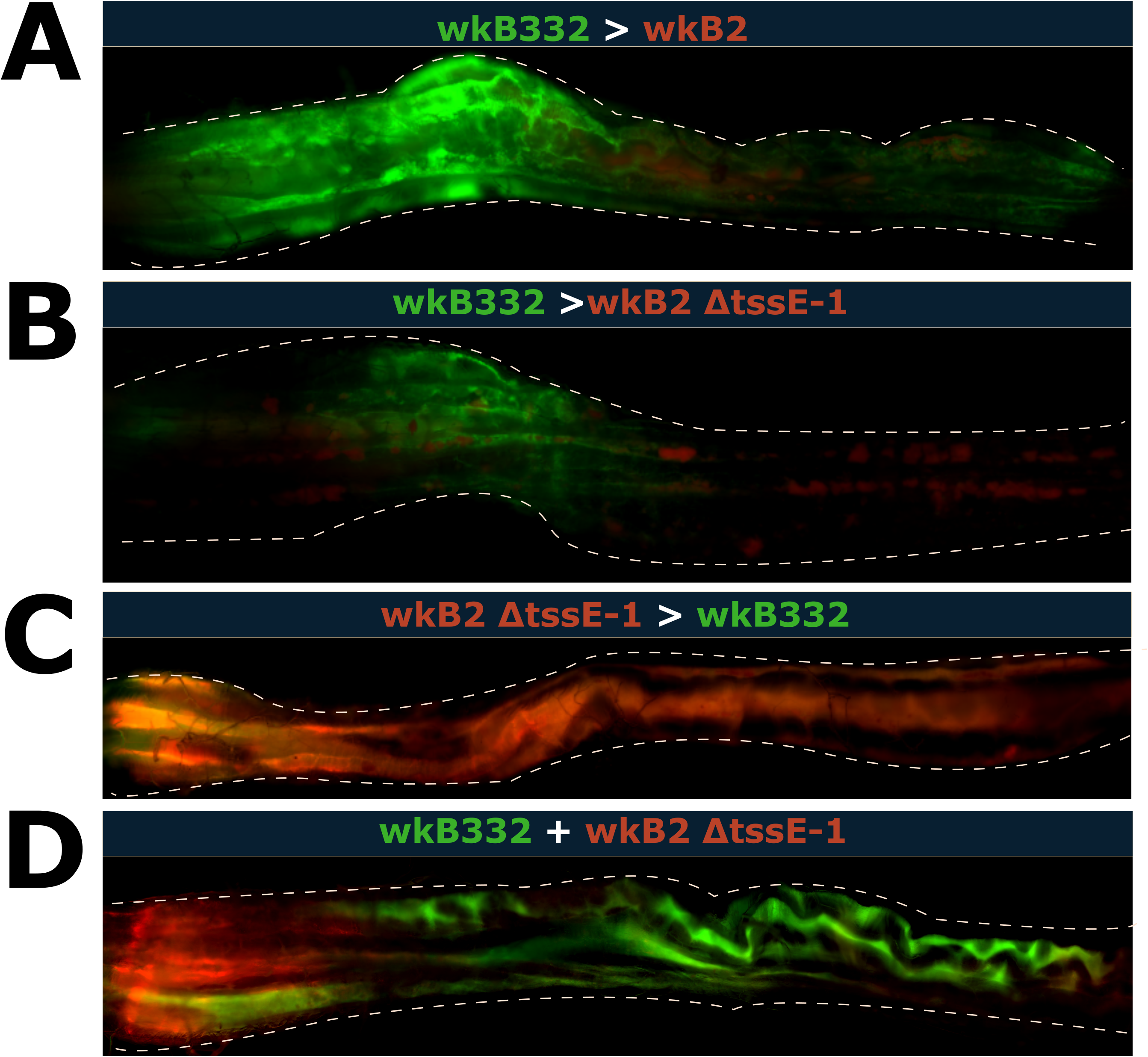
Images of honey bee gut colonization treatments in which co-existence was observed, generated through fluorescent microscopy. The image includes only the ileum section of the gut and is oriented from pylorus-adjacent to rectum-adjacent, left to right. Wild type wkB332 is colored green in these images, while wild type wkB2 and wkB2 ΔtssE-1 are colored red. **A)** wkB332 followed by wkB2, **B)** wkB332 followed by wkB2 ΔtssE-1, **C)** wkB2 ΔtssE-1 followed by wkB332, **D)** wkB332 simultaneously inoculated with wkB2 ΔtssE-1.

## Discussion

Microbiome composition reflects the interplay between stochastic and deterministic factors. Here, we demonstrate the strengths of the honey bee gut microbiome as a system for understanding that interplay. Additionally, this work provides support for priority effects as a mechanism by which weakly competitive symbiont strains might persist in the presence of more competitive strains. The relationship between *S. alvi* strains wkB2 and wkB332 is reaffirmed through our experiments, confirming that the T6SS-1 system of wkB2 contributes to its ability to routinely outcompete wkB332 in directly competitive scenarios [31, 35]. We build on this understanding by showing that temporal advantages can support the persistence of wkB332 in the presence of wkB2, with the strength of these advantages dependent on the presence of the T6SS.

Priority effects are thought to affect compositional outcomes across systems in community ecology. The term has been used as early as 1987 to describe the ability of dominant plant species to prevent the establishment of late-emerging competitors [51]; conversely, early arriving taxa may facilitate subsequent colonists [52]. In the case of intraspecific competition between strains of bacteria, the early arriving taxa can completely or partially exclude subsequent taxa [18, 53]. In our experimental design, we utilized two lengths of delayed inoculations to better understand how time might influence these outcomes. For our two isolates, we saw consistent inhibition of invaders at our longer delay period of five days; however, only wkB2 consistently repelled invasion in our one-day delay experiments. Honey bees establish their gut microbiomes by approximately the fourth day post-emergence [43]. Our results suggest that, after this period of establishment, strain proportions of *S. alvi* within the gut become relatively stable.

Because honey bees do not leave the hive until ∼21 days post-emergence [54], the inability of late-arriving strains to colonize might exclude *S. alvi* strains encountered outside the hive, including *Snodgrassella* strains associated with other bee species foraging at the same flowers. Thus, the delay in exposure may contribute to the host specificity documented for *Snodgrassella* [55, 56], although host-specific regulatory mechanisms appear also to contribute [57, 58].

In our one-day delay experiments, we saw distinct outcomes based on strain identity and the presence or absence of a working T6SS-1 system. When the T6SS-1 possessing strain wkB2 was inoculated first, wkB332 was greatly inhibited. Disabling the T6SS-1 increased the ability of wkB332 to invade, indicating that the T6SS-1 helps to exclude later arriving *S. alvi* strains. The *S. alvi* T6SS-1 system was also beneficial when invading an established *S. alvi* population, as *S. alvi* wkB2 was more able to invade wkB332 when the T6SS-1 was active. Other mechanisms affect competitive success in these strains, as even the wkB2 knockout achieves a higher relative abundance than wkB332, regardless of order of arrival.

Because the T6SS action is contact-dependent, the advantage it confers to the more abundant strain is confined to its immediate surroundings. Potentially, different strains, each with distinct toxin-immunity systems, could establish at different locations in the uncolonized gut and ultimately dominate within local patches. If so, an established community within a single bee gut might contain multiple strains, each confined to distinct regions. Colonization patterns of fluorescently tagged strains gave some evidence of such patchiness; however, our images of intact guts do not resolve fine-scale spatial organization. Depending on whether the mechanism of antagonism is localized, as for T6SS, or more dispersed, as for diffusible toxins such as bacteriocins [59], different outcomes might be expected.

Overall, our results provide evidence of priority effects in the establishment of the honeybee microbiome. Potentially, these effects help to explain the persistence of multiple strains of *S. alvi* in populations of honey bees [34]. We found that the magnitude of priority effects depends on the timing of arrival of different strains, which, in the context of the honey bee life cycle, may help to enforce host specificity. We confirm the previously established idea that competitive outcomes between T6SS-possessing and vulnerable strains operate in a deterministic fashion [31, 35]. Our findings show that stochasticity in which strain colonizes first can impact these outcomes and can mitigate T6SS-derived advantages. This study joins a growing body of research providing experimental evidence of priority effects within microbial communities and prompts further exploration into the interplay of deterministic and stochastic processes in determining microbiome composition in hosts.

## Acknowledgments

Funding support was provided by a Stengl-Wyer Postdoctoral Scholarship to KRJ, a National Institutes of Health award (R35GM131738) to NAM, and a National Science Foundation award (NSF IOS 2103208) to NAM and Jeffrey Barrick. We thank Erick Motta, PJ Lariviere, E. Powell, Jeffrey Barrick, and Alan Cerqueira for advice on the study, and K. Hammond and E. Powell for logistical support including apicultural assistance.

## References

1. Moossavi S et al. The prebiotic and probiotic properties of human milk: implications for infant immune development and pediatric asthma. Front Pediatr 2018;6. 10.3389/fped.2018.00197

2. Rebollar EA et al. Integrating the role of antifungal bacteria into skin symbiotic communities of three Neotropical frog species. ISME J 2019;13:1763. 10.1038/s41396-019-0388-x

3. Sommer F, Bäckhed F. The gut microbiota — masters of host development and physiology. Nat Rev Microbiol 2013;11:227–238. 10.1038/nrmicro2974

4. Zheng H et al. Metabolism of toxic sugars by strains of the bee gut symbiont Gilliamella apicola. mBio 2016;7:e01326–16. 10.1128/mBio.01326-16

5. Hild B et al. Neonatal exposure to a wild-derived microbiome protects mice against diet-induced obesity. Nat Metab 2021;3:1042–1057. 10.1038/s42255-021-00439-y

6. Woodhams DC et al. Prodigiosin, violacein, and volatile organic compounds produced by widespread cutaneous bacteria of amphibians can inhibit two Batrachochytrium fungal pathogens. Microb Ecol 2018;75:1049–1062. 10.1007/s00248-017-1095-7

7. Powell JE et al. Field-realistic tylosin exposure impacts honey bee microbiota and pathogen susceptibility, which is ameliorated by native gut probiotics. Microbiol Spectr 2021;9:10.1128/spectrum.00103-21. 10.1128/spectrum.00103-21

8. Kohl KD et al. Microbial detoxification in the gut of a specialist avian herbivore, the Greater Sage-Grouse. FEMS Microbiol Lett 2016;363. 10.1093/femsle/fnw144

9. Liberti J et al. Gut microbiota influences onset of foraging-related behavior but not physiological hallmarks of division of labor in honeybees. mBio 2024;15:e01034–24. 10.1128/mbio.01034-24

10. Kramp RD, Kohl KD, Stephenson JF. Skin bacterial microbiome diversity predicts lower activity levels in female, but not male, guppies, Poecilia reticulata. Biol Lett 2022;18:20220167. 10.1098/rsbl.2022.0167

11. Burz SD et al. From microbiome composition to functional engineering, one step at a time. Microbiol Mol Biol Rev 2023;87:e00063–23. 10.1128/mmbr.00063-23

12. Debray R et al. Priority effects in microbiome assembly. Nat Rev Microbiol 2021. 10.1038/s41579-021-00604-w

13. Vellend M. Conceptual Synthesis in Community Ecology. Quart Rev Biol 2010;85:183–206. 10.1086/652373

14. Zha Y et al. Different roles of environmental selection, dispersal, and drift in the assembly of intestinal microbial communities of freshwater fish with and without a stomach. Front Ecol Evol 2020;8:152. 10.3389/fevo.2020.00152

15. Jones KR, Hughey MC, Belden LK. Colonization order of bacterial isolates on treefrog embryos impacts microbiome structure in tadpoles. Proc R Soc B 2023. 10.1098/rspb.2023.0308

16. Delory BM et al. When history matters: The overlooked role of priority effects in grassland overyielding. Funct Ecol 2019;33:2369–2380. 10.1111/1365-2435.13455

17. Jones KR, Belden LK, Hughey MC. Priority effects alter microbiome composition and increase abundance of probiotic taxa in treefrog tadpoles. Appl Environ Microbiol 2024;90:e00619–24. 10.1128/aem.00619-24

18. Devevey G et al. First arrived takes all: inhibitory priority effects dominate competition between co-infecting Borrelia burgdorferi strains. BMC Microbiology 2015;15:61. 10.1186/s12866-015-0381-0

19. Chen JZ et al. A strong priority effect in the assembly of a specialized insect-microbe symbiosis. Appl Environ Microbiol 2024;0:e00818–24. 10.1128/aem.00818-24

20. Fukami T. Historical contingency in community assembly: integrating niches, species pools, and priority effects. Ann Rev Ecol Evol Syst 2015;46:1–23. 10.1146/annurev-ecolsys-110411-160340

21. Chappell CR et al. Wide-ranging consequences of priority effects governed by an overarching factor. eLife 2022;11:e79647. 10.7554/eLife.79647

22. Fukami T, Mordecai EA, Ostling A. A framework for priority effects. J Veget Sci 2016;27:655–657. 10.1111/jvs.12434

23. Zhou J, Ning D. Stochastic community assembly: Does it matter in microbial ecology? Microbiol Mol Biol Rev 2017;81. 10.1128/MMBR.00002-17

24. Fargione J, Brown CS, Tilman D. Community assembly and invasion: An experimental test of neutral versus niche processes. Proc Natl Acad Sci USA 2003;100:8916–8920. 10.1073/pnas.1033107100

25. Anderson MC et al. Shigella sonnei encodes a functional t6ss used for interbacterial competition and niche occupancy. Cell Host Microbe 2017;21:769-776.e3. 10.1016/j.chom.2017.05.004

26. Granato ET, Meiller-Legrand TA, Foster KR. The evolution and ecology of bacterial warfare. Curr Biol 2019;29:R521–R537. 10.1016/j.cub.2019.04.024

27. Russell AB et al. Type VI secretion delivers bacteriolytic effectors to target cells. Nature 2011;475:343–347. 10.1038/nature10244

28. Jani AJ, Cotter PA. Type VI secretion: Not just for pathogenesis anymore. Cell Host Microbe 2010;8:2–6. 10.1016/j.chom.2010.06.012

29. Coulthurst S 2019. The Type VI secretion system: a versatile bacterial weapon. Microbiology ;165:503–515. 10.1099/mic.0.000789

30. Ma L-S et al. Agrobacterium tumefaciens deploys a superfamily of type VI secretion DNase effectors as weapons for interbacterial competition in planta. Cell Host Microbe 2014;16:94–104. 10.1016/j.chom.2014.06.002

31. Speare L et al. Bacterial symbionts use a type VI secretion system to eliminate competitors in their natural host. Proc Natl Acad Sci USA 2018;115:E8528–E8537. 10.1073/pnas.1808302115

32. Zheng H et al. Honey bees as models for gut microbiota research. Lab Anim (NY) 2018;47:317–325. 10.1038/s41684-018-0173-x

33. Kwong WK, Moran NA. Gut microbial communities of social bees. Nat Rev Microbiol 2016;14:374–384. 10.1038/nrmicro.2016.43

34. Ellegaard KM, Engel P. Genomic diversity landscape of the honey bee gut microbiota. Nat Commun 2019;10:446. 10.1038/s41467-019-08303-0

35. Motta EVS et al. Type VI secretion systems promote intraspecific competition and host interactions in a bee gut symbiont. Proc Natl Acad Sci USA 2024;121:e2414882121. 10.1073/pnas.2414882121

36. Martinson VG et al. A simple and distinctive microbiota associated with honey bees and bumble bees. Mol Ecol 2011;20:619–628. 10.1111/j.1365-294X.2010.04959.x

37. Martinson VG, Moy J, Moran NA. Establishment of characteristic gut bacteria during development of the honeybee worker. Appl Environ Microbiol 2012;78:2830–2840. 10.1128/AEM.07810-11

38. Bobay L-M, Wissel EF, Raymann K. Strain structure and dynamics revealed by targeted deep sequencing of the honey bee gut microbiome. mSphere 2020;5:e00694–20. 10.1128/mSphere.00694-20

39. Steele MI et al. Diversification of type VI secretion system toxins reveals ancient antagonism among bee gut microbes. mBio 2017;8:e01630–17. 10.1128/mBio.01630-17

40. Steele MI, Moran NA. Evolution of interbacterial antagonism in bee gut microbiota reflects host and symbiont diversification. mSystems 2021;6:e00063–21. 10.1128/mSystems.00063-21

41. Horak RD, Leonard SP, Moran NA. Symbionts shape host innate immunity in honeybees. Proc R Soc B 2020;287:20201184. 10.1098/rspb.2020.1184

42. Elston KM et al. The Pathfinder plasmid toolkit for genetically engineering newly isolated bacteria enables the study of Drosophila-colonizing Orbaceae. ISME Commun 2023;3:1–11. 10.1038/s43705-023-00255-3

43. Powell JE et al. Routes of acquisition of the gut microbiota of the honey bee Apis mellifera. Appl Environ Microbiol 2014;80:7378–7387. 10.1128/AEM.01861-14

44. R Core Team. R: A language and environment for statistical computing. 2021. Vienna, Austria: R Foundation for Statistical Computing, 2021.

45. RStudio Team. RStudio: Integrated development for R. 2020. Boston, MA: RStudio, PBC, 2020.

46. Benjamini Y, Hochberg Y. Controlling the false discovery rate: a practical and powerful approach to multiple testing. J R Stat Soc B Methodol 1995;57:289–300.

47. Wickham H. ggplot2: Elegant graphics for data analysis. 2016. Springer-Verlag New York, 2016.

48. Alboukadel K. ggpubr: ‘ggplot2’ based publication ready plots. 2023. 2023.

49. Hadley W, Pederson TL, Seidel D. scales: Scale functions for visualization. 2023. 2023. https://scales.r-lib.org/reference/scales-package.html

50. McMurdie PJ, Holmes S. phyloseq: An R package for reproducible interactive analysis and graphics of microbiome census data. PLoS ONE 2013;8:e61217. 10.1371/journal.pone.0061217

51. Quinn JF, Robinson GR. The effects of experimental subdivision on flowering plant diversity in a california annual grassland. J Ecol 1987;75:837–855. 10.2307/2260209

52. Gurung A et al. Strain-dependent and host genotype–dependent priority effects in gut microbiome assembly affect host fitness in. Limnol Oceanogr 2024;69:1782–1796. 10.1002/lno.12614

53. Segura Munoz RR et al. Experimental evaluation of ecological principles to understand and modulate the outcome of bacterial strain competition in gut microbiomes. ISME J 2022;16:1594–1604. 10.1038/s41396-022-01208-9

54. Abou-Shaara HF. The foraging behaviour of honey bees, Apis mellifera: a review. Vet Med 2014;59:1–10. 10.17221/7240-VETMED

55. Cornet L et al. Phylogenomic analyses of Snodgrassella isolates from honeybees and bumblebees reveal taxonomic and functional diversity. mSystems 2022;7:e01500–21. 10.1128/msystems.01500-21

56. Li Y et al. Species divergence in gut-restricted bacteria of social bees. Proc Natl Acad Sci USA 2022;119:e2115013119. 10.1073/pnas.2115013119

57. Guo L et al. Reactive oxygen species are regulated by immune deficiency and Toll pathways in determining the host specificity of honeybee gut bacteria. Proc Natl Acad Sci USA 2023;120:e2219634120. 10.1073/pnas.2219634120

58. Kwong WK et al. Genomics and host specialization of honey bee and bumble bee gut symbionts. Proc Natl Acad Sci USA 2014;111:11509–11514. 10.1073/pnas.1405838111

59. Heilbronner S et al. The microbiome-shaping roles of bacteriocins. Nat Rev Microbiol 2021;19:726–739. 10.1038/s41579-021-00569-w

